# Microbiota metabolic exchange is critical for colorectal cancer redox homeostasis and growth

**DOI:** 10.1101/2020.12.02.408013

**Authors:** Hannah Bell, Joshua Goyert, Samuel A. Kerk, Nupur K. Das, Costas A. Lyssiotis, Yatrik M. Shah

## Abstract

Intestinal microbiota play a fundamental role in human health and disease. Microbial dysbiosis is a hallmark of colorectal cancer (CRC) as tumor stage-specific shifts potentiate tumor growth, influence the inflammatory microenvironment, and alter response to therapy. Recent work has demonstrated a critical role for microbial metabolite exchange in host response. However, the role of most microbial metabolites in colon cancer growth is unclear. To better understand how metabolic exchange between the microbiota and tumor epithelium alter CRC growth, a screen of the most abundant bacterially derived metabolites was assessed. Several metabolites were found to alter CRC growth, but reuterin most significantly suppressed CRC cell proliferation. Reuterin is a bifunctional metabolite containing both hydroxy and aldehyde functional groups. Reuterin is primarily synthesized from glycerol by *Lactobacillus reuteri,* a commensal bacterium found throughout the gastrointestinal tract. We found that reuterin suppresses growth via alterations to the redox balance of CRC cells. Mechanistically, reuterin potentiates reactive oxygen species (ROS) which leads to irreversible cysteine oxidation and enhanced cell death. Supplementation of either antioxidants or hydrogen sulfide fully rescued growth, suggesting that reuterin is suppressing CRC growth through protein oxidation. These studies demonstrate the potential of reuterin to act as a potent chemotherapeutic for treating colorectal cancers.

## Introduction

CRC is the third-most common cancer and accounts for 10% of all cancer-related deaths (1). The global rate of CRC has increased 40% since 1975 and is projected to increase by 60% in the next 15 years. An estimated 13 million people are estimated to die from colon cancer in the year 2030 (1). Unhealthy eating habits, low physical activity, and obesity are major environmental risk factors believed to be responsible for the rising rate of CRC development (2). There are several known genetic predispositions that significantly increase a patient’s risk for CRC including Lynch syndrome and familial adenomatous polyposis (3). The development of CRC typically follows a uniform progression from a polyp that develops into an adenoma which can progress into colorectal cancer (3). The tumors interface with a diverse and highly dense population of microbes collectively termed the microbiota. The typical human gastrointestinal tract contains more than 10^14^ microbial organisms that are predominately composed of 500-1000 unique bacterial species, but also includes archaea, protists, and fungi (4). An intact microbiome is crucial for health in humans.

In healthy patients, bacteria that colonize the gastrointestinal tract develop a symbiotic relationship with their host. The microbiota breaks down otherwise indigestible compounds, secretes metabolites, aids in development of the mucosal layer, and prevents detrimental pathogens from colonizing in the gut (5). Short-chain fatty acids (SCFAs), like propionic acid and butyric acid, are a well-characterized class of bacterial-derived metabolite that regulate inflammation and act as an energy source for the intestinal epithelium of their host (6). Bacterial species of the gut can also secrete essential vitamins for host consumption, such as vitamin K, vitamin B2, vitamin B12, and folates (7). The metabolites produced by the microbiota are sensed by both non-hematopoietic and hematopoietic cells, which stimulates the development of the mucosal and humoral immune systems (8). In a murine model, germ-free animals have significantly less antibody production and defective gut-associated lymphoid tissue (9). Additionally, germ-free mice develop significantly more and larger colorectal tumors, relative to conventionally raised mice in mouse models cancer model (10). The microbiota has a protective role against colorectal cancer, but disturbances in the resident communities can create a state of pathological dysbiosis.

The relationship between dysbiosis and CRC is poorly characterized. Although incomplete, current evidence strongly suggests that CRC is characterized by increased levels of the pathogenic species *Fusiobacteria, Heliobacteria, Acinetobacter*, and *Psuedomona* (16). A comparative analysis of microbiota found that adenomas correlated with a more significant shift in distal gut microbiota than diet, body mass index, or familial history of CRC. Additionally, a combined metagenomic and metabolic analysis found that there are significant stage-specific shifts within the microbiota of patients with polypoid adenomas and intramucosal carcinomas (17). Collectively, these studies highlight the ability of colorectal cancers to alter the composition of their surrounding microbiota. The present work aims to better characterize the dynamic relationship between the microbiota and colorectal tumors by assessing how metabolites generated by bacterial species influence CRC proliferation and progression. A screen of bacterial-derived metabolites found that several microbial compounds reduce CRC growth. Reuterin was the most potent and at physiological relevant concentrations led to cell death in colon cancer cell lines. RNA-sequencing and metabolomic analysis revealed that reuterin significantly alters cellular redox balance. Mechanistically, we demonstrate that reuterin increases cysteine oxidation leading to cell death. These findings suggest that maintaining the microbial metabolite exchange during colon tumorigenesis can tumor selective protein oxidation and a decrease in colon cancer proliferation and progression. Metabolites produced by the microbiota are currently an untapped resource of potential pharmacological agents that could offer significant therapeutic benefits to future cancer patients.

## Materials and Methods

### Cell culture

HCT116, DLD1 and SW480 cells were maintained at 37°C in 5% CO_2_ and 21% O_2_. Cells were cultured in Dulbecco’s Modified Eagle Medium (DMEM) supplemented with 10% FBS and 1% antibiotic/antimycotic.

### MTT Assay

2000 cells were plated into a 96-well cell culture plate in 200 μl of supplemented DMEM. 72 hours after treatment, cells were incubated with 50 μl of 3-(4,5-dimethylthiazolyl-2)-2, 5-diphenyltetrazolium bromide (MTT) dye for 45 minutes at 37° Celsius and 5% carbon dioxide in a cell-culture incubator. Media and dye were aspirated following the incubation period and the remaining formazan was dissolved in 75 μl of dimethyl sulfoxide for 5 minutes. The absorption was then measured at 570 nm. The readings taken at 72 hours were normalized to the day 0 values for data analysis.

### Colony Forming Assay

500 cells were plated in a 6 well plate, allowed to grow for three days before treatment with indicated compound. After 7 additional days of growth cells were washed once with PBS, fixed for 5 minutes in Formalin, then stained for 5 minutes with Crystal Violet and destained with water.

### Cell death assay

25,000 cells were plated into 500 μl of supplemented DMEM in a 24-well cell culture plate and allowed to adhere for 24 hours prior to experimental treatment. Following experimental treatment, the media was pipetted into a sterile 1.5 ml Eppendorf tube and 125 μl of trypsin was added to each well for 3 minutes. Following the 3-minute incubation period, the original media was added back into its respective well and cells were pipetted off of the cell culture plate. 30 μl of each well was added into a sterile 1.5 ml Eppendorf tube and mixed with 30 μl of 0.4% trypan blue dye. 10 μl of this mixture was counted on a hemocytometer to quantify cell death.

### RT-qPCR

300,000 cells were plated in biological triplicates in 6-well cell culture plates in 2 mL of supplemented DMEM. Cells were left to adhere to the plate for 24 hours prior to experimental treatments. Following experimental treatment, the media and treatments were aspirated from each well and 1 mL of trizol was administered to each well. The 1 mL of trizol was transferred to a sterile 1.5 mL Eppendorf tube and then 200 μl of chloroform was added. The samples were vortexed for 15 seconds and set to incubate for two minutes. The samples were then centrifuged for 15 minutes at 16.1 × 1000 relative centrifugal force (rcf) at 4°C. 350 μl of the top aqueous layer was removed and added to a sterile 1.5 mL Eppendorf tube. 500 μl of isopropanol was added to each sample and vortexed. Following an incubation period of 15 minutes on ice, each sample was centrifuged at 16.1 rcf and 4 degrees Celsius for 15 minutes. Following centrifugation, the supernatant was aspirated and 1 mL of 75% ethanol was added to each sample. Each sample was then centrifuged for 10 minutes at 16.1 rcf at 4 degrees Celsius. The ethanol supernatant was then aspirated and the purified RNA isolate was dissolved in 30 μl of sterile nuclease-free water. cDNA was generated using the ThermoFisher High-capacity cDNA Reverse Transcription Kit. The product from reverse transcription was diluted in 80 μl of sterile nuclease free water. qPCR was run in a 384-well plate. Every sample was loaded in three technical triplicates for each gene of interest. Each well was loaded with 5 μl of qPCR SYBR green master mix, 3.4 μl of sterile nuclease-free water, 0.3 μl of the forward primer (**Table 1**), .3 μl of the reverse primer (**Table 1**), and 1 μl of the diluted cDNA product.

### Enteroid Culture

Colon was isolated from CDX2-ER^T2^CreApc^fl/fl^ mice and cut into 1cm pieces. Tissue was incubated in 10mM DTT for 15 minutes at room temperature. Tissues were rinsed with DPBS supplemented with gentamicin and primocin. Tissue was incubated with slow rotation at 4°C for 75minutes in 8mM EDTA. EDTA was removed and tissue was put through three cycles of snap-shakes to release crypts. Isolated crypts were spun down and collected in cold LWRN medium. Crypts were then plated in matrigel (Corning) in 96-well culture plates in LWRN media.

### Whole genome RNA-sequencing and analysis

Healthy intestinal *Rattus norvegicus-*derived IEC6 cells were treated with 150 μM of reuterin for 24 hours prior to analysis. RNA sequencing libraries were prepared using the TruSeq RNA library prep kit v2 (Illumina, San Diego, CA) following the manufacturer’s recommended protocol. The libraries were sequenced using single-end 50-cycle reads on a HiSeq 2500 sequencer (Illumina) at the University of Michigan DNA Sequencing Core Facility. RNA-sequencing quality control of raw fastq files was performed using FastQC v 0.11.5. Fastq files were mapped to the rat genome using STAR-2.5.3.a using the options “outFilterMultimapNmax 10” and “sjdbScore 2”. Gene expression levels were quantified using Subread v1.5.2 package FeatureCounts. Differential expression testing was conducted with the Bioconductor package edgeR v3.16.5 using glmLRT. To reduce the dispersion of the dataset due to lowly expressed genes, genes with a mean aligned read count less than five across all samples were excluded from analysis. Genes were considered differentially expressed as indicated, either at a false discovery rate (FDR) of < 0.01 or < 0.1 to yield high and low stringency approaches.

### ROS Assay

The cell-permeable free radical sensor carboxy-H2DCFDA (Invitrogen) was used to measure intracellular ROS levels. Cells were harvested by ice-cold PBS-EDTA(5mM) buffer and incubated with 10 μM carboxy-H2DCFDA in PBS at 37°C for 20 min. The cells were washed, resuspended in PBS and analyzed using Beckman Coulter MoFlo Astrios flow cytometer. Data was analyzed using FlowJo software. Values are expressed as the percentage of cells positive for DCF fluorescence.

### Lactate Dehydrogenase Leakage Assay

The cell death after reuterin treatment in various cell lines was evaluated using an LDH Cytotoxicity Detection Kit. Briefly, cells were exposed to various concentrations of reuterin or rescue compound for 24 h. 100 μL of cell-free media was transferred in triplicate to a new 96 well plate, then 100 uL of the LDH reaction mixture was added. After 30 minutes of incubation in the dark at room temperature, the optical density of the final solution was determined at a wavelength of 490 nm using a microplate reader.

### Statistical Analysis

Data represent the mean ± s.e.m or s.d. in case of viability experiments. Data are from three independent experiments measured in triplicate, unless otherwise stated in the figure legend. For statistical analyses, Student’s *t*-tests were conducted to assess the differences between two groups. One-way ANOVA was used for multiple treatment conditions. A *P* value of less than 0.05 was considered to be statistically significant. All statistical tests were carried out using Prism 8 software (GraphPad).

## Results

### Microbiota-derived metabolite screen identifies novel compounds with significant inhibitory effects on CRC cell growth

To determine the effect of intestinal commensal-derived metabolites on CRC proliferation, a cell growth assay was conducted in colon cancer-derived HCT116 cells. 42 exclusively bacterial derived metabolites identified in the colon were assessed for their impact on cell growth (**Figure 1a,b**). Of this expansive list, pyridoxal hydrochloride, tyramine, 1-3 diaminopropane, and reuterin were found to have the most robust inhibitory effects on growth in HCT116 cells. Selected metabolites with potential for growth repression were then tested at lower doses in HCT116 cells (**Figure 1c)**. Dose-response curves in three colon cancer-derived cell lines identified that reuterin possessed the greatest inhibitory growth effect relative to all other screened metabolites. (**Figure 1d).** A reuterin dose curve was done in eight additional cancer cell lines demonstrating consistently robust growth inhibition (**Figure 2a-c**). Moreover, decreases in growth were observed at concentrations that are physiologically relevant and observed in the intestine (18). The decrease in cell growth was attributed to cell death as assessed by trypan blue and LDH assays (**Figures 2d and e**). Moreover, reuterin is in equilibrium with the unsaturated aldehyde acrolein, which has been shown to be cytotoxic to cells (19). Consistent with this data acrolein was cytotoxic to several colon cancer cell lines (**Figure 2f).** To determine if primary tumor cells are sensitive to reuterin, we generated tumor enteroids derived from an *Apc* deleted colon adenoma mouse model (20). Reuterin led to cell death and decreased enteroid growth in this model (**Figure 2g).** Together this data demonstrates the endogenous microbial metabolite reuterin can potently decrease colon cancer growth.

**Figure 1:**
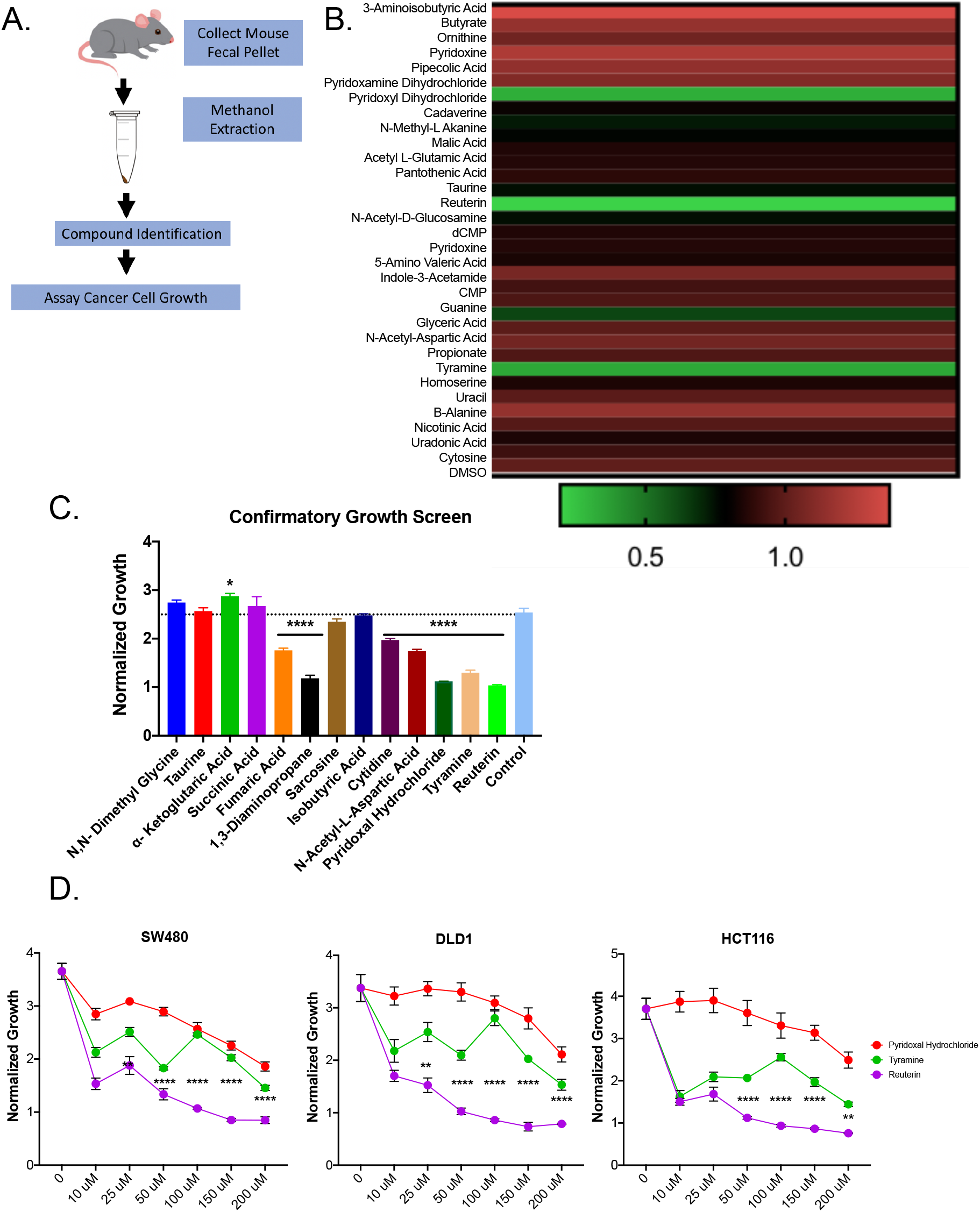
A screen of microbial derived compounds identified several that reduced colon cancer cell growth in vitro. A. Schematic diagram of the screen. B. 42 compounds were incubated with HCT116 cells for 3 days at 1mM and growth was assessed by MTT. C. Selected compounds were also tested on HCT116 cells for 3 days at 500μM. D. Each cell line was treated with a range of doses of pyridoxyl hydrochloride, tyramine, and reuterin and assessed via MTT assay.

**Figure 2:**
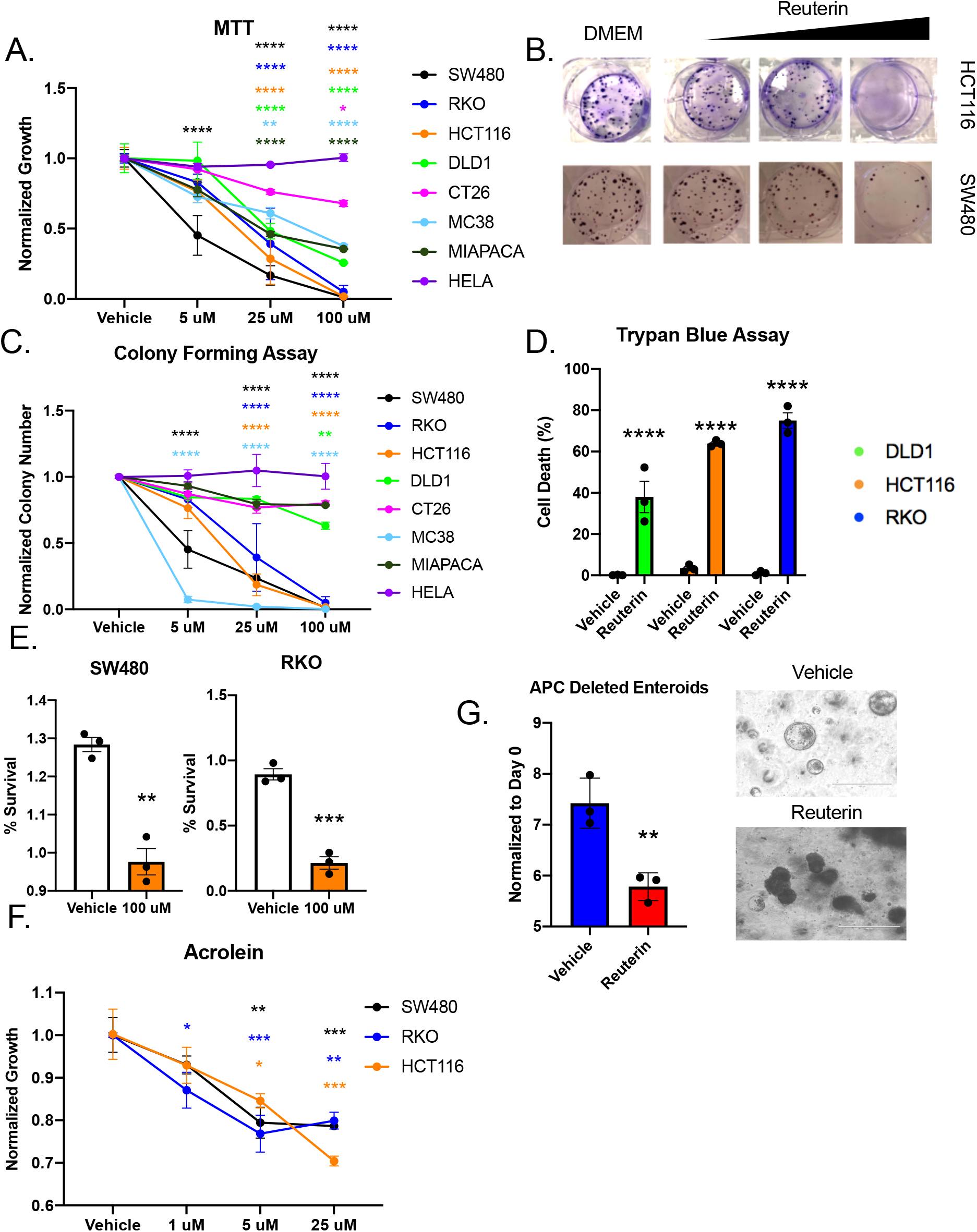
Reuterin inhibits proliferation and induces cell death in cancer cell lines. A. Cell lines were assessed by MTT for proliferation at indicated doses of reuterin after 3 days. B. Image and C. quantification of colony forming assay cancer cell lines. Cell death assessed by D. trypan blue or E. LDH release. F. MTT following treatment with acrolein. G. Growth of colorectal cancer mouse enteroids after reuterin treatment.

### Reuterin dysregulates redox balance to inhibit CRC cell growth

To begin to elucidate the mechanism behind the decrease in cell growth following reuterin treatment, whole-genome mRNA analysis was assessed via RNA-seq. Reuterin treatment led to significant changes in the transcription of 1,255 genes (**Figure 3a**). Gene set enrichment showed particularly striking changes in serine glycine biosynthesis, the pentose phosphate pathway, and integrin signaling (**Figure 3b).** Further analysis of the RNA-seq data identified that several antioxidant NRF2 target genes were significantly increased in the presence of reuterin (**Figure 3c**) (21). NRF2 levels are low in most cells via degradation initiated by Kelch-like ECH-associated protein 1 (KEAP1) (22). Under oxidative stress, cysteine modification on KEAP1 dissociates the KEAP1-NRF2 complex, allowing stabilization and activation of NRF2 can directly activate NRF2 by dissociating KEAP1 (22, 23). Maintaining proper redox balance is essential for cancer cell growth. Targeted metabolomics in HCT116 and SW480 cell lines confirmed altered glutathione metabolism (**Figure 3d).** We observed a significant increase in oxidized L-Glutathione after 24 hours following reuterin treatment in both HCT116 and SW480 cells (**Figure 3e)**. The RNA-seq and metabolomics data suggest an increase oxidative stress.

**Figure 3:**
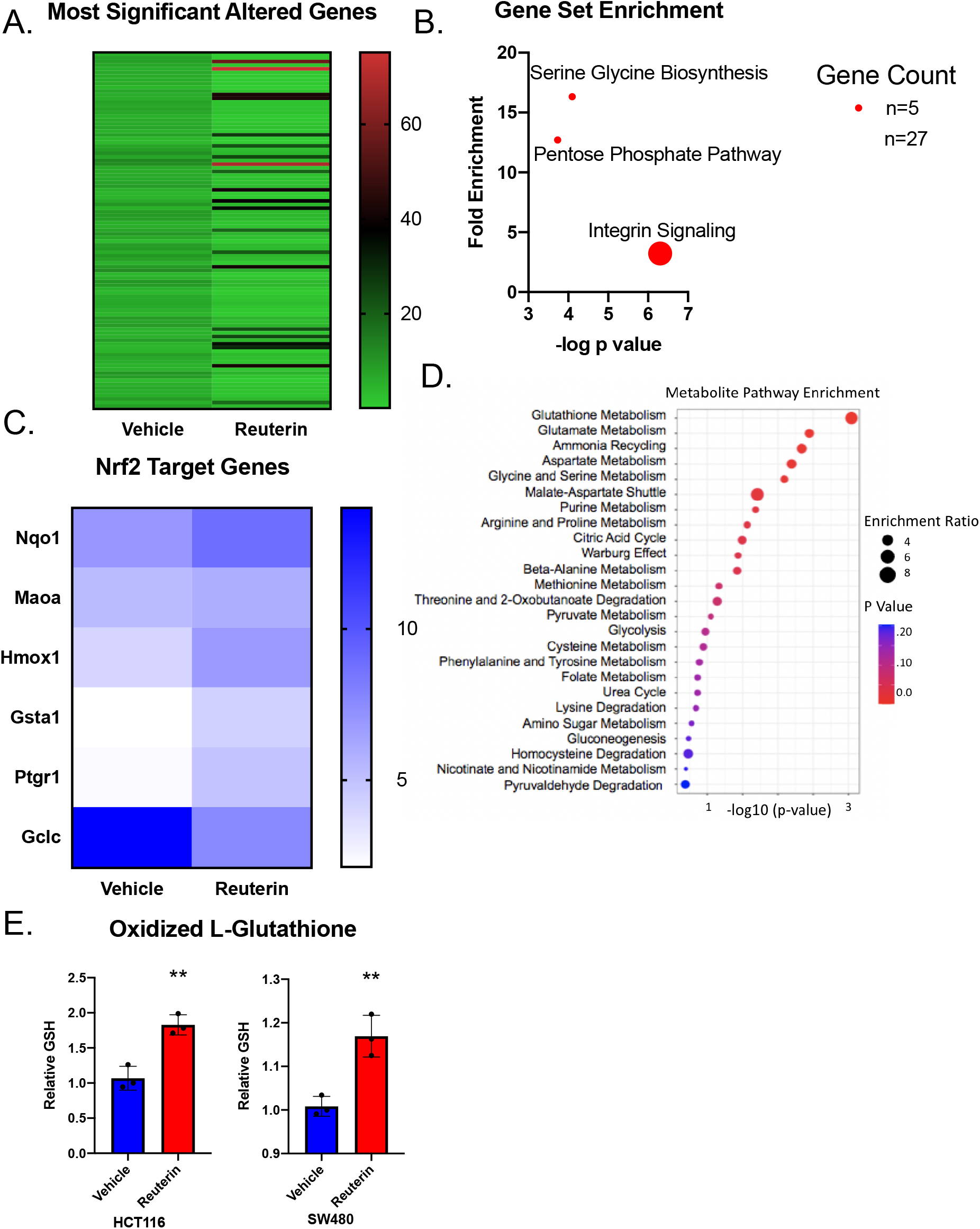
Combined RNA-SEQ and metabolomics data suggests reuterin induces oxidative stress in colorectal cancer cells. A. Cells were treated with reuterin for 24 hours and RNA was collected for RNA-SEQ. B. Enriched pathways after PANTHER KEGG analysis. C. Heatmap of Nrf2 target genes identified in the RNA-seq analysis. D. Pathway enrichment analysis of metabolomics data after colon cancer cells were treated with reuterin for 24 hours. E. Quantification of increase of oxidized glutathione after 24-hour treatment reuterin in HCT116 and SW480 cell lines.

### Reuterin activates the Nrf2 pathway in CRC Cells

Reuterin resulted in a significant increase in ROS in multiple cell lines (**Figure 4a)**. To confirm NRF2 activation, qPCR analysis was performed on NRF2 and several well characterized NRF2 target genes including HMOX1, GCLM, and NQO1 in HCT116 and SW480. The data demonstrated that reuterin activates the NRF2 pathway in a dose-dependent manner (**Figure 4b**).

**Figure 4:**
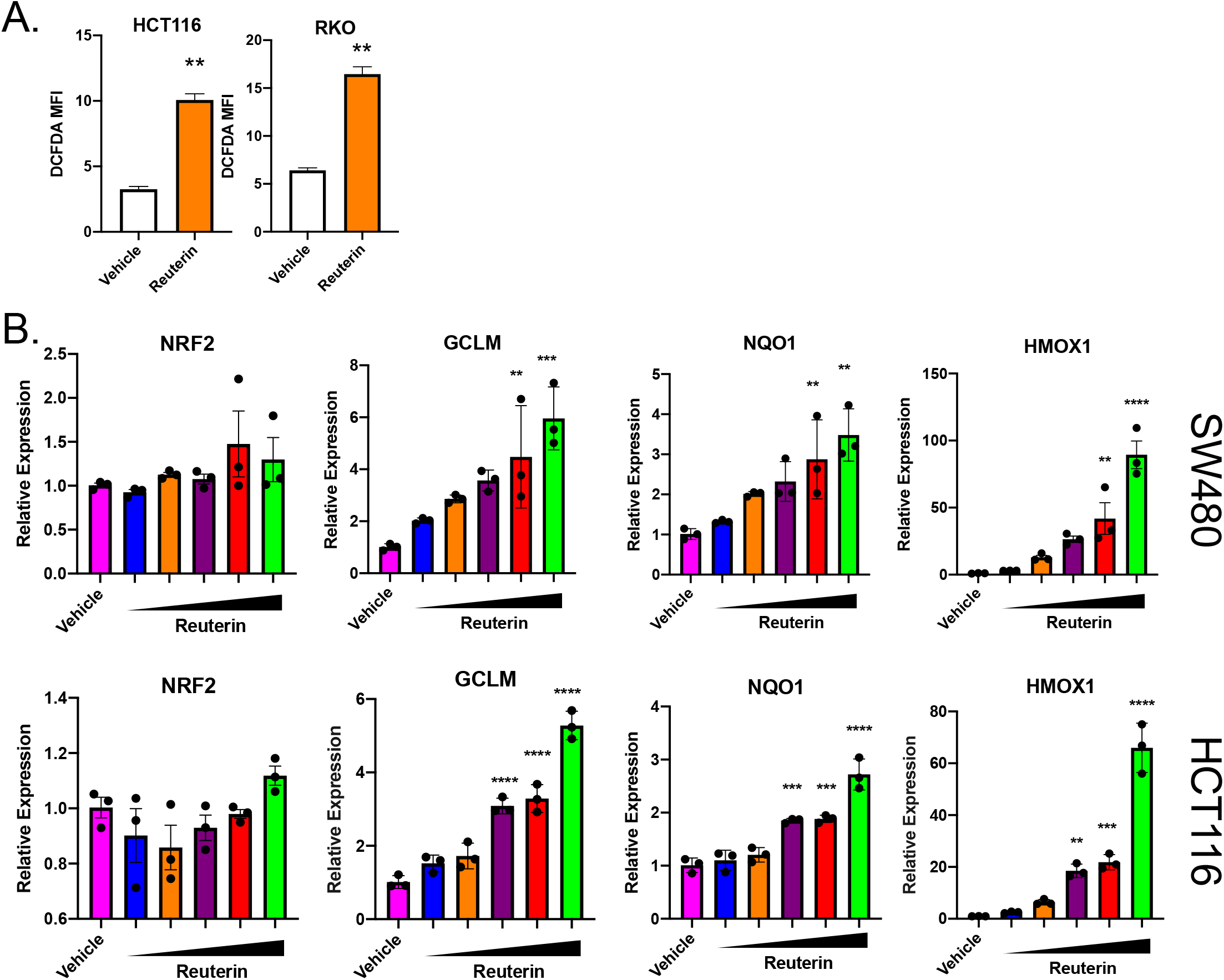
Reuterin increases ROS and Nrf2-target genes. A. Cells were treated with indicated concentration for 24 hours then stained with DCFDA dye and acquired. B. Nrf2 target gene activation was assessed after 24 hours of incubation with increasing reuterin doses from 5 to 150 uM.

To assess if heightened oxidative stress results in cell death following reuterin treatment, an MTT assay was performed in HCT116, SW480, and DLD1 cells that were co-treated with reuterin and N-acetyl cysteine (NAC). Reuterin treatment inhibited cell growth and, in a dose dependent manner NAC rescued cell death (**Figure 5a and b**). We also confirmed that NAC rescues reuterin-induced gene expression changes (**Figure 5c).** Mechanistically, NAC replenishes glutathione concentrations. Glutathione is a potent antioxidant that can inhibit free radicals, peroxides, lipid peroxides, and heavy metals (24). To understand the selectivity of ROS generated by reuterin, two additional antioxidants, liproxstatin-1 and ferrostatin-1, were assessed (**data not shown**). Liproxstatin-1 and ferrostatin-1 detoxify lipid- and metal-catalyzed radicals (25). Liproxstatin-1 or ferrostatin-1 were not able rescue growth and their inability to protect against reuterin treatment suggests that reuterin may selectively enhance free radicals. Hydrogen sulfide can protect enhanced oxidation of cysteine thiols via persulfidation, a posttranslational modification (**Figure 6a**, 26). Sodium sulfide was also able to rescue reuterin-induced cell death (**Figure 6b).** This demonstrates that reuterin is inducing protein oxidation and this is a major mechanism leading to cancer cell death.

**Figure 5:**
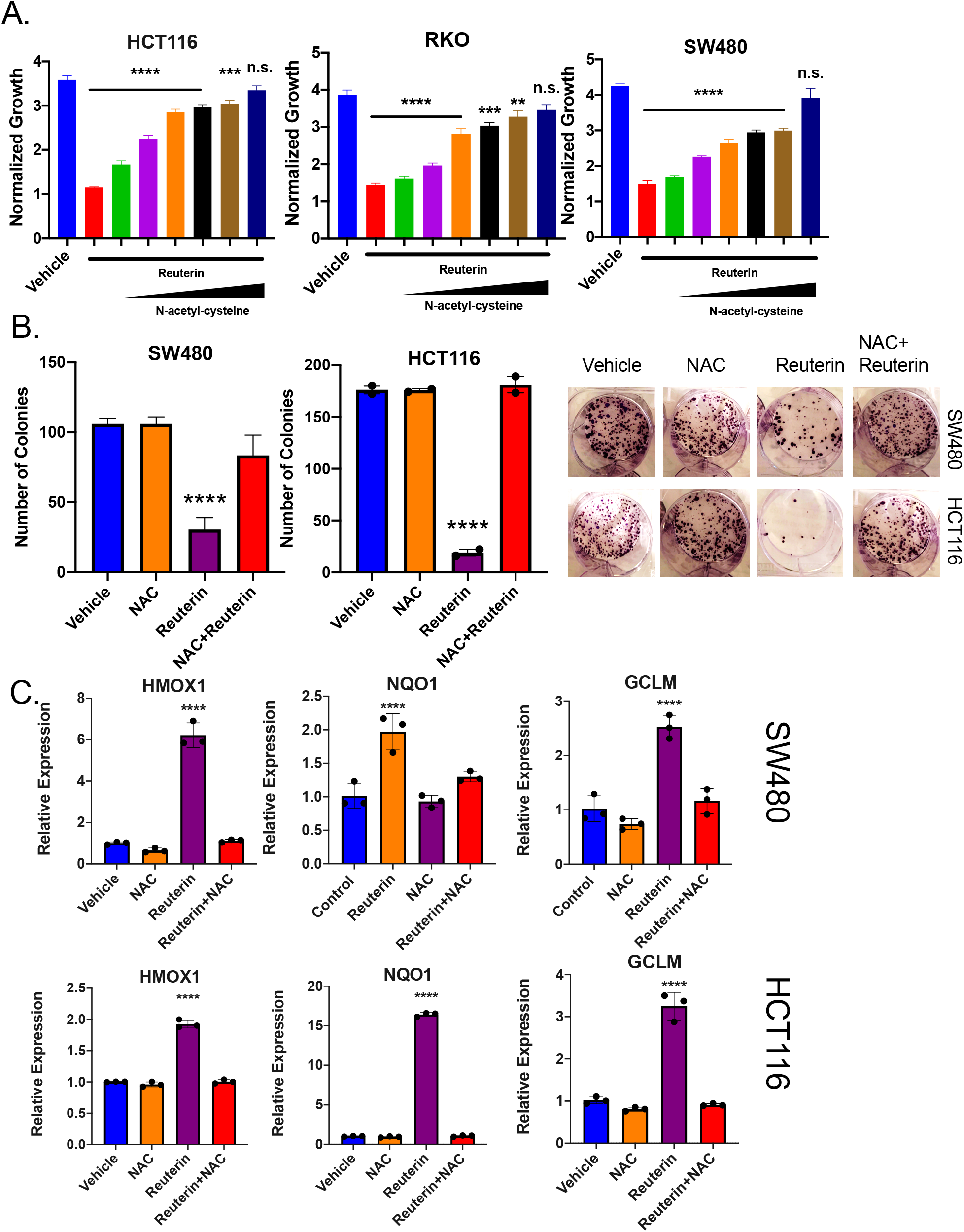
NAC protects from reuterin-induced cell death. A. MTT and B. colony forming assays and C. gene expression analysis of cells pretreated with increasing doses of NAC then treated with reuterin for 3 days.

**Figure 6:**
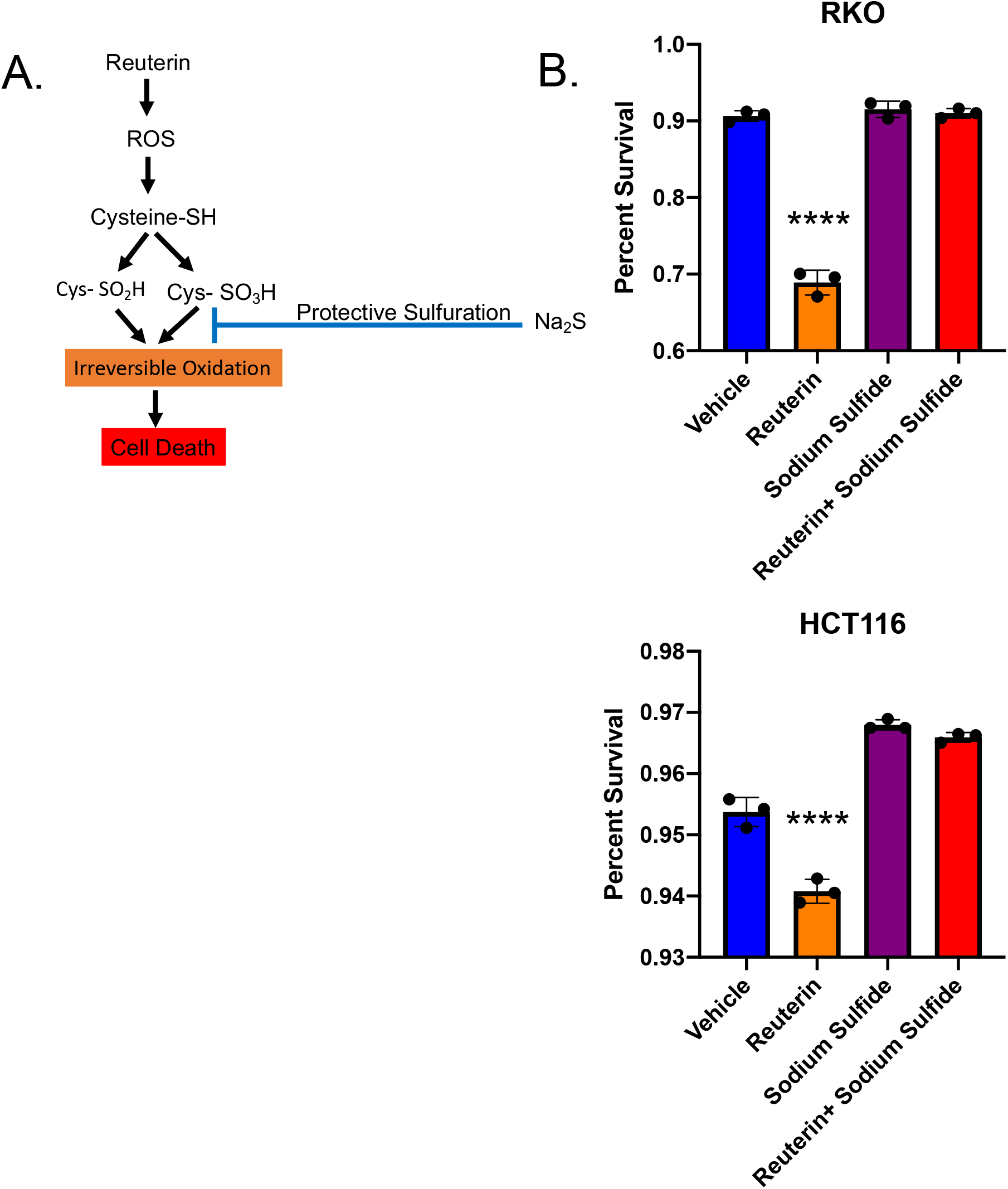
Sodium Sulfide protects from reuterin induced cell death. A. Schematic of sodium sulfide protection. B. LDH release in cells pretreated with sodium sulfide for 6 hours, then treated with reuterin for 24 hours.

## Discussion

The gut microbiota impacts many facets of our physiology and dysbiosis can lead to diseases of the intestine, such as inflammatory bowel disease (27). Moreover, the intestinal microbiome can have far reaching effects that influence metabolic diseases, neurological disorders, and cognitive function (28, 29). Recent work has highlighted the intimate relationship between intestinal microbiota and CRC progression. *Fusobacterium nucleatum* increased CRC proliferation rates in vitro and tumor burden in murine-cancer models. Alternatively, several commensal species of bacteria have shown promise by reducing tumor formation in murine-cancer models (30, 31). The role of specific bacterial species in promoting or inhibiting colorectal cancer is poorly characterized, but current literature demonstrates there is a strong connection between CRC and the microbiome.

Chemotherapeutics and immunotherapies are largely ineffective without an intact gut microbiota (32). Tumors that were once thought to arise from sterile sites have shown local microbiota changes that are essential for their growth (33). Advanced 16s rRNA sequencing and metagenomic analyses enable researchers to thoroughly investigate the species present within the tumor microenvironment. However, the causal role and mechanistic insights into the role of the microbiota in tumorigenesis remains unclear. Metabolomic data of colon contents suggest that the microbiota produces a repertoire of unique metabolites that are poorly understood with respect to the impact on the tumor microenvironment.

The relationship between bacterial-derived metabolites and CRC is an emerging area of research. Butyrate, one of the most abundant metabolites induced cell death in colorectal, gastric, and breast cancer by inhibiting histone deacetylase (HDAC) (34). HDACs remove acetyl groups and lysine residues from histones which allows DNA to form transcriptionally silenced chromatin. Inhibition of HDACs block this process resulting in altered gene expression that can ultimately lead to cell death (35). The beneficial impacts of butyrate are typically seen at 10 mM concentrations in vitro and 50 mM in vivo (34). The low potency of this compound makes these laboratory findings difficult to apply in a pharmacological context. Moreover, several more potent HDAC inhibitors are FDA approved and are currently undergoing clinical trials for multiple cancers (35). Our research aimed to identify more potent metabolites that impact on CRC growth at lower concentrations. Through a functional screen of 42 gut-microbial metabolites, the present work identified several compounds that are robust inhibitors of CRC growth at micromolar doses. The metabolites examined throughout our investigation are far more potent inhibitors of proliferation than butyrate.

Reuterin, also known as 3-hydroxypropionaldehyde, has a simple molecular structure that consists of a 3-carbon backbone with an aldehyde and alcohol functional group on carbons 1 and 3 respectively (36). Reuterin was identified in our microbial metabolite screen as a potent suppressor of colon cancer cell growth. Interestingly, work from our group has shown that reuterin is a potent inhibitor of hypoxia-inducible factor (HIF)-2α, a central transcription factor in CRC growth and progression, suggesting additional mechanisms of action for reuterin (37). The administration of reuterin to CRC cells induces the antioxidant NRF2 pathway, which is typically activated in the presence of heightened oxidative stress. The NRF2 activation and cell death following reuterin treatment is completely attenuated upon co-treatment with the antioxidant NAC. This suggests that the functional downstream pathway of reuterin is due to heightened oxidative stress, which we verified through direct ROS measurement. Reuterin induces ROS in *Escherichia coli,* but little is known about its mechanism in eukaryotic cells (38). Aldehydes and alcohols are known to generate ROS within prokaryotic and eukaryotic organisms, and we aim to determine if the aldehyde moiety in reuterin is sufficient to activate ROS in cancer cells (39).

Redox homeostasis is essential for tumor growth and survival. Reactive oxygen species are highly reactive oxygen molecules. ROS form via oxidative cellular metabolism and at low levels are important mediators of cell signaling and function (22, 23). High levels of ROS disrupt redox homeostasis and are highly toxic to cancer cells. Due to their highly altered metabolism, microenvironmental inflammatory response, and rapid proliferation, cancers have unique mechanisms to tightly regulate ROS formation and redox balance (22, 23). Antioxidants have historically been used to reduce tumorigenesis in target cancers. However, a battery of new data suggests this approach promotes cancer growth and survival (40). As an alternative, an increase in ROS concentrations to toxic levels in cancer cells is an efficacious mechanism for treatment (41). Many known chemotherapeutics increase ROS, which cause irreparable damage or death, in cancers. However, this approach has deleterious side effects on nearby cells. Due to the paucity of healthy colon cell lines, we were not able to directly assess the selectivity of reuterin in colon cancer cells compared to normal colon cells, but reuterin has significant inhibitory effects on proliferation at a concentration of 10 μM in cancer-derived cell lines. The physiological concentration of reuterin in the colon is typically 100 μM, and healthy intestinal cells continue to proliferate in its presence, suggesting there is significant pharmacological potential for this compound. Interestingly, reuterin’s inhibition of growth appears to be fairly colon specific, as pancreatic and cervical cancer cell lines were less sensitive to reuterin

Our future experiments will aim to elucidate the specific mechanism behind the inhibitory effect of reuterin on colorectal cancer growth. Additionally, we will seek to clarify if the oxidative stress induced by reuterin is conserved among other cancers. To further assess the clinical relevance of reuterin as a chemotherapeutic, we will test if probiotic use of *Lactobacillus reuteri* has protective effects in colon cancer.

## Notes

**Conflict of Interests**: The authors declare no conflict of interest

### Competing Interest Statement

The authors have declared no competing interest.

